# Self-buckling of undulating flagella: an elastohydrodynamic mechanism for double waves in spermatozoa

**DOI:** 10.64898/2026.09.11.750800

**Authors:** Pyae Hein Htet, Kenta Ishimoto

**Affiliations:** Department of Mathematics, Kyoto University, Kyoto 606-8502, Japan

## Abstract

The relatively long flagella of spermatozoa from insects, birds, and octopuses display double waves, characterized by two superimposed helical waves. The prevalance of these highly organized waveforms across diverse taxa and distinct flagellar architectures hints at shared underlying physics, motivating a model of the flagellum as an elastic filament immersed in a viscous fluid, actively driven by a single set of internal bending moment waves. Simulations of a clamped filament show that it can buckle under its own activity into whirling and flapping states. A multiple-scales analysis of the elastohydrodynamic equations reveals how nonlinear interactions between fast undulations generate an effective compression driving buckling, and connects wave-driven buckling to classical follower-force instabilities. Extending the model to a swimming spermatozoon, the same instability produces double waves. Parameter estimates across species show that most observed double waves lie within the regime where buckling is permitted, supporting self-buckling as a generic physical mechanism for double waves.

## I. INTRODUCTION

The spermatozoon plays a fundamental role at the beginning of development. Motility generated by the beating of its flagellum enables it to navigate the fluid environment towards the egg, leading to fertilization and the development of a new organism. A large body of research spanning multiple fields has therefore been devoted to understanding the flagellar beat, from its ultrastructural anatomy and internal processes generating oscillations [1], to the fluid-structure interactions converting flagellar waves into locomotion [2].

Much of our current understanding of sperm motility comes from mammalian spermatozoa and model organisms such as sea urchins and zebrafish. However, spermatozoa across the wider biological world display a far richer variety of beating patterns and flagellar architectures. A notable example is the double-wave, or superhelical, waveform (Fig. 1), characterized by short-wavelength, small-amplitude bending waves, referred to as minor waves, superimposed on a slower, larger-scale, helical major wave.

**FIG. 1.**
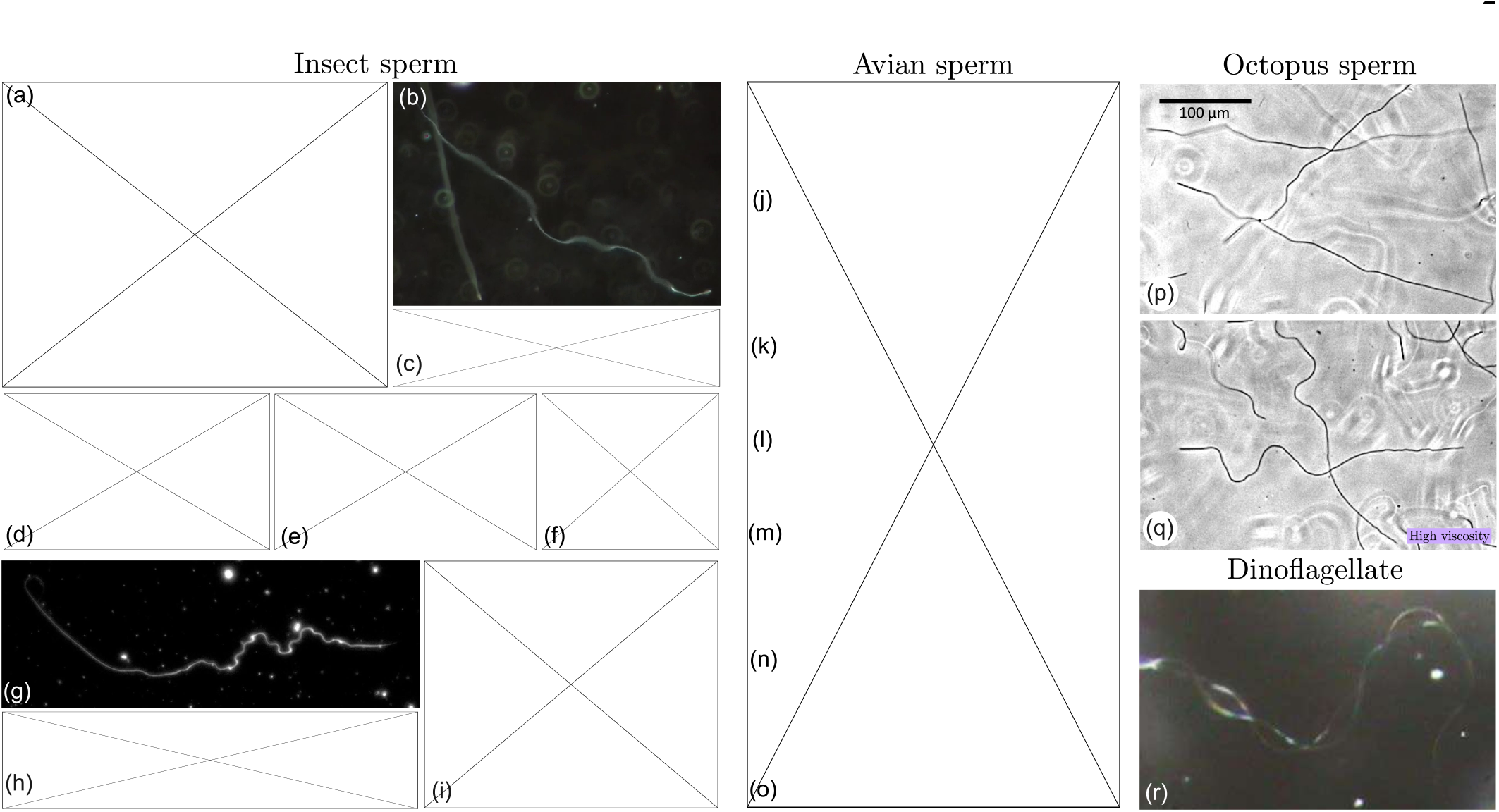
Double waves across the biological world. (a-i) Double waves in insect spermatozoa: (a) *Culex quinquefasciatus* [13] (b) *Anopheles gambiae* [14] (c) *Megaselia scalaris* [5] (d) *Bacillus rossius* [3] (e) *Tenebrio molitor* [4] (f) *Aleochara curtala* [9] (g) *Sitotroga cerealella* [15] (h) *Culicoides melleus* [6] (i) *Aedes notoscriptus* [12]. (j-q) Avian spermatozoa exhibiting single waves in a water-like medium and double waves in a high-viscosity medium (1.5 Pa s methylcellulose solution) [16]: (j-k) *Gallus domestica* (l-m) *Coturnix japonica* (n-o) *Columba livia*. (p-q) *Octopus vulgaris* spermatozoa exhibiting single waves in a water-like medium and double waves in a 1.5 Pa s methylcellulose solution. (r) Double waves in an unidentified dinoflagellate [17]. Images reproduced with permission (b) from *Nyasembe et al*. [14]. Copyright © 2025 The Authors, CC BY-NC-ND 4.0; (g) from Shah *et al*. [15]. Copyright © 2023 The Authors, CC BY 4.0; (p-q) courtesy of Yuuko Wada; (r) from Cosson & Prokopchuk [17], with permission from Galina Prokopchuk. Some figures are excluded from the preprint due to copyright restrictions and may be accessed from the respective references.

Since its discovery in the 1970s in the stick insect [3] and the mealworm beetle [4], the double-wave morphology has been documented in widely separated insect groups, including flies [5–8], beetles [9–11], mosquitoes [12–14] and moths [15], as well as in avian spermatozoa [16], and more recently, in octopus sperm and even dinoflagellates [17]. Despite the remarkable taxonomic diversity of double waves, illustrated in Fig. 1, it remains poorly understood how these highly organized waveforms emerge from the underlying flagellar activity.

An important feature common to these flagella is the motile core, known as the axoneme, which typically consists of nine outer microtubule doublets arranged around a central pair of singlet microtubules, forming the canonical 9+2 structure [1]. Dynein motors anchored to the outer doublets consume ATP to generate active shear forces between neighbouring doublets. Structural constraints, imbued by nexin links between adjacent outer doublets and by radial spokes extending inward towards the central pair, convert dynein-induced sliding into bending [18, 19]. The coordinated activity of dynein motors along the axoneme thus drives the bending waves responsible for cell motility [20].

Beyond this common basic plan, however, flagella exhibit enormous ultrastructural variation across taxa. The sea urchin sperm flagellum is among the simplest, consisting essentially of a plasma membrane around a 9+2 axoneme. Passerine avian spermatozoa are distinguished by nine outer dense fibres surrounding the axoneme [21], a feature common also to mammalian [22] and octopus sperm [23]. Insect spermatozoa are some of the most complex, consisting of nine additional singlet microtubules around a 9+2 structure [24] (“9+9+2”), although the 9+9+1 architecture of mosquito sperm [25] presents a notable exception. Many insect sperm also have proteinaceous accessory bodies and elongated mitochondrial derivatives alongside the axoneme [24].

For decades, double waves were documented almost exclusively in insect spermatozoa, mainly alongside detailed ultrastructural studies, and the hypotheses in these early studies, that the major and minor waves are generated by distinct structural components of the insect flagellum [3, 4, 12], remains the prevailing view despite limited direct evidence. Ref. [9] proposed an alternative explanation that only the minor waves are actively propagated along an intrinsically helical flagellum, while the major waves are an illusion created by the rolling of the cell.

However, the recurrence of the double-wave morphology across such diverse taxa and flagellar anatomy suggests a possible origin not rooted in particular biological features specific to certain spermatozoa, but rather, in the physics common to actively beating flagella. This is further supported by experiments examining the motility of the same spermatozoa in different fluid environments, in which double waves were seen exclusively in [16] or strongly promoted by high-viscosity media [5], pointing to a crucial role of the spermatozoon’s mechanical environment.

Together with the conserved axonemal core, this motivates a description of the flagellum as an internally driven elastic rod interacting with the surrounding viscous fluid, known as an elastohydrodynamic model. Here, the coordinated activity of molecular motors is coarse-grained into an effective continuous forcing, and the mechanical behavior of the axoneme and its accessory structures is represented through effective material properties.

Many foundational two-dimensional elastohydrodynamic studies have investigated the emergence and regulation of the flagellar beat [20, 26–29], but a three-dimensional formulation is required to describe more complex waveforms. Here the flagellum is represented as a Kirchhoff rod [30], in which a material frame attached to the centerline tracks bending and twisting deformations, and its motion is coupled to the surrounding viscous fluid using a hydrodynamic theory of choice [31–33]. However, these equations are often severely numerically stiff and computationally demanding, motivating the more efficient asymptotic coarse-grained approach of Ref. [34] which has subsequently been extended from planar filaments to three dimensions [35]. The resulting dramatic improvements in efficiency have made extensive simulations of strongly nonlinear three-dimensional filament dynamics feasible.

Alongside these computational developments, several studies have revealed the importance of elastohydrodynamic instabilities in flagellar dynamics. Ref. [36] showed, within a planar model, that travelling waves of internally generated shear can induce a buckling instability which breaks waveform symmetry and enables curved swimming trajectories.

Recent three-dimensional simulations showed that non-planar beats can spontaneously emerge from planar internal moments, producing waveforms resembling those observed in bull and sea urchin sperm [37].

A complementary line of research considers follower force models, in which the internal activity is represented by an imposed tangential force. Ref. [38] used a detailed two-filament flagellar model to show that steady dynein activity can induce a dynamic buckling instability and sustain beating without invoking a dynein regulation mechanism. Ref. [39] showed that a simpler model, consisting of a follower force applied at the free end of an elastic filament, produces a similar instability. Although this model was developed in the context of a molecular motor translocating along a cytoskeletal filament, it closely resembles the mechanical setting of a flagellum, especially when generalized to accommodate distributed forces along the centerline [40], and has become a fundamental model for the dynamics of active biological filaments.

In our study, we apply these theoretical and computational developments to the long-standing question of how the superhelical waveform arises. We hypothesize that the major wave emerges as an instability of a filament subject to a single set of ‘minor’ bending waves, without the need for separate generation of major and minor waves. After isolating the instability in the simpler setting of a clamped filament, we show that the same buckling instability indeed reproduces double waves in a swimming spermatozoon. A multiple-scales mathematical analysis accompanying our computational results not only rationalizes them, but also reveals a unifying description of wave-driven buckling and classical follower force instabilities. A more detailed theoretical analysis predicts the range of mechanical parameters within which buckling is possible, and parameter estimates for numerous species collated from the literature allow us to assess the relevance of this mechanism across biologically diverse spermatozoa.

The paper is organized as follows. In §II, we formulate the mathematical model of an internally driven elastic filament in a viscous fluid. We present numerical results for a clamped filament in §III and analyze the instability theoretically in §IV. We then extend the numerical and theoretical analyses to a swimming spermatozoon in §V and §VI, respectively. In §VII, we evaluate our results in the broader biological context, before concluding in §VIII.

## II. MODEL SETUP

The conserved axonemal architecture motivates a mathematical model of the flagellum as an actively driven elastic filament in a viscous fluid (see Fig. 2a). We consider a slender, inextensible filament of length *L*, immersed in a Newtonian fluid of viscosity *µ*, modelled as a homogeneous Kirchhoff rod with a circular cross-section of radius *ϵL* and bending rigidity *EI*. Parametrizing the centerline by **r**(*s, t*), where *s* is arclength and *t* is time, the unit tangent vector is given by **t** = **r**^*′*^(*s, t*), henceforth using ′ and · to denote derivatives with respect to *s* and *t* respectively.

**FIG. 2.**
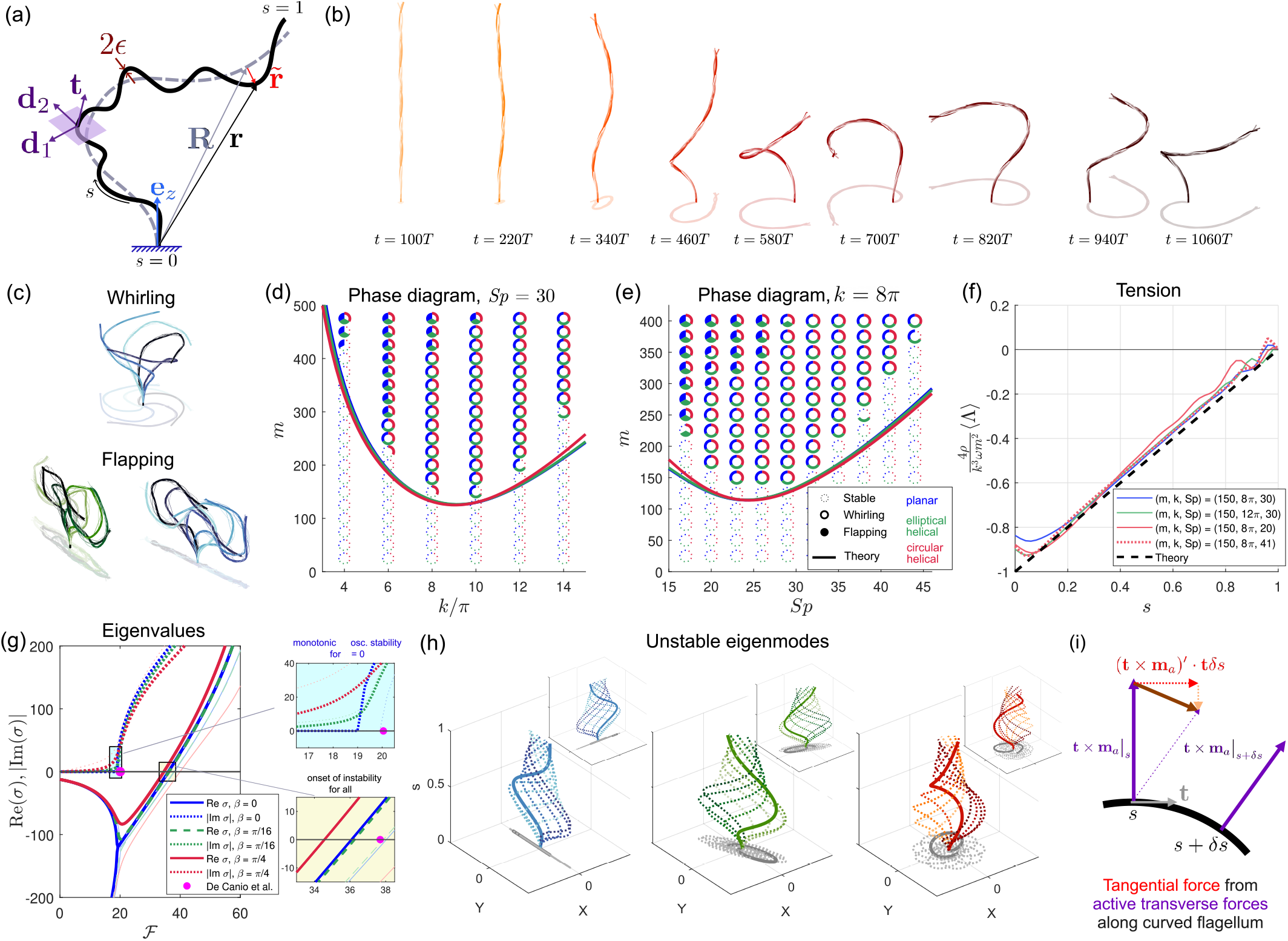
Numerical and theoretical results for a clamped filament. (a) Schematic of the problem setup, a clamped elastic filament in a viscous fluid subject to active internal moments. The multiple-scales expansion is also illustrated. (b) Time sequence of a clamped filament buckling into a whirling state. Filament snapshots from the preceding period are shown as thin lines to illustrate minor waves. Parameters used are (*Sp, k, m, β*) = (30, 8*π*, 250, *π/*4). (c-d) Filament snapshots at different times superposed to illustrate the whirling (c) and flapping states (d). Parameters used are (*Sp, k*) = (30, 8*π*), with *m* = 250 for whirling and *m* = 425 for flapping. Time progresses from light to dark color. (d-e) Phase diagram illustrating long-time filament behavior in (*m, k, Sp*) parameter space for fixed *Sp* = 30 (d) and fixed *k* = 8*π*. (e). Unfilled markers with dotted outlines indicate the stable straight time-averaged state. Unfilled markers with solid outline indicates whirling, and filled markers indicate flapping. *β* = 0, *π/*16, *π/*4 are indicated by blue, green, and red, respectively. Solid lines plot the buckling threshold *m*_*c*_ from linear stability analysis of the slow equation. (f) Time-averaged tension profiles (normalized) computed numerically for various parameter values (colored lines) plotted together with the theoretically calculated tension profile (black dashed). (g) Real (solid/dashed) and imaginary (dotted) parts of the most unstable eigenvalue branch plotted in thick line against 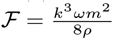, for parameters (*Sp, k*) = (30, 8*π*). The next most unstable eigenvalue branches are plotted as thinner lines with lighter color. Insets zoom in to the monotonic-oscillatory stable transition and the onset of stability. (h) The corresponding leading (main panel) and sub-leading (inset) eigenmodes. Blue/green/red color scheme is used for *β* = 0, *π/*16, *π/*4, respectively. (i) Schematic showing tangential force (red) arising from transverse active forces (purple) on an infinitesimal section of a curved filament. Their combined effect is mean compression.

### A. Elasticity

The internal elastic force and moment are denoted by **F**(*s, t*) and **M**(*s, t*) respectively. At microscopic scales, fluid and filament inertia are negligible. The equations of elasticity consist of the force balance **F**^*′*^ + **f**_*h*_ = **0**, the moment balance **M**^*′*^ + **t** × **F** + **m**_*a*_ + **m**_*h*_ = **0**, and the Euler-Bernoulli constitutive relationship [41, 42] **M** = *EI***t** × **t**^*′*^ + **M**_3_ relating the elastic moment to the filament’s geometric and material properties. Here **f**_*h*_ and **m**_*h*_ are the hydrodynamic force and moment densities, respectively, and **m**_*a*_ is the active internal moment density. **M**_3_ is the contribution from filament twist, specified fully in SI Appendix. We define the tension Λ := **F** · **t**.

### B. Hydrodynamics

Elasticity is coupled to the surrounding fluid by specifying **f**_*h*_ and ***m***_*h*_ (full details in SI Appendix) using a local drag approximation, known in fluid mechanics as ‘resistive-force theory.’ In this framework, **f**_*h*_ is anisotropically proportional to local filament velocity, with different drag coefficients *c*_⊥_ = 2*c*_∥_ = 4*πµ/*[log(2*/ϵ*) − 0.5] [43] reflecting the greater hydrodynamic resistance to perpendicular than tangential filament motion. Unless stated otherwise, we fix *ϵ* = 0.001, relevant to many insect and avian spermatozoa. A similar local drag theory relates fluid torque to angular velocity about the tangent with a rotational drag coefficient *c*_*r*_ = −4*πµϵ*^2^*L*^2^ [44].

### C. Internal activity

Internal activity is prescribed as a travelling wave of active bending moment density **m**_*a*_(*s, t*) = **m** exp(*ik*_∗_*s* − *iω*_∗_*t*), with wavenumber *k*_∗_ and frequency *ω*_∗_. The amplitude **m** lies in the plane normal to the local tangent, spanned by the basis vectors **d**_1_(*s, t*) and **d**_2_(*s, t*). We write **m** = *m*_1_**d**_1_ + *im*_2_**d**_2_, with *m*_1_, *m*_2_ ∈ ℝ. Up to a basis rotation, this is the most general active moment amplitude with constant **d**_1_ and **d**_2_ components.

Writing *m* = |**m**| and *β* = arctan(−*m*_2_*/m*_1_), we may equivalently write **m** = *m*(cos *β***d**_1_ −*i* sin *β***d**_2_). We will henceforth focus on *β* = *π/*4, *π/*16 and 0. The circular helical active moment *β* = *π/*4 is relevant to spermatozoa exhibiting helical waves, whereas *β* = 0 corresponds to purely planar forcing, an idealization of planar beating. Real flagellar beats, however, are not perfectly planar, and we represent a small out-of-plane component of activity with an elliptical helical forcing *β* = *π/*16. Values of *β* outside [0, *π/*4] straightforwardly map into this range by an appropriate rotation or reflection.

### D. Governing equations for subsequent multiple-scales analysis

The preceding equations are numerically solved using the scheme in Ref. [35], as outlined in the Methods. They may also be manipulated (as detailed in SI Appendix) to yield coupled equations for the filament centerline **r** and tension Λ. Nondimensionalizing lengths by *L*, forces by *EI/L*^2^, and time by the elastohydrodynamic relaxation timescale *c*_⊥_*L*^4^*/EI*, these equations may be written as

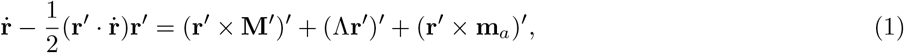

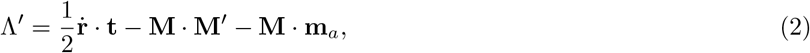

with **M** = **t** × **t**^*′*^, neglecting twist contributions for subsequent theoretical analysis (as well justified by the insensitivity of simulation results to twist effects). The dynamics are governed by the dimensionless groups: the dimensionless wavenumber *k* = *k*_∗_*L* and the sperm number

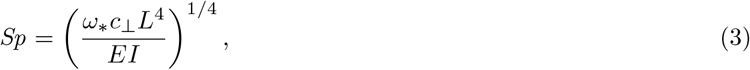

which quantifies the relative importance of viscous resistance to elastic relaxation. The dimensionless frequency of internal activity is *ω* = *Sp*^4^, so that *Sp* has entered our system through the active moment, **m**_*a*_ = **m** exp (*iks* − *iωt*).

In water-like media, mammalian sperm typically have *Sp <* 10, whereas longer insect sperm exhibiting double waves can reach *Sp* ∼ 50; the 2 mm-long *Drosophila melanogaster* sperm [7] is an extreme case with *Sp* ∼ 250. Since *Sp* ∝ *µ*^1*/*4^, a 1500-fold increase in viscosity raises *Sp* by around a factor of 6, so that superhelical avian spermatozoa in such media can reach *Sp* ∼ 100.

## III. NUMERICAL SIMULATIONS OF A CLAMPED FILAMENT

We first investigate elastohydrodynamic instabilities in the simple setting of a filament clamped at one end and free at the other. We therefore impose the boundary conditions **r** = **0, r**^*′*^ = **e**_*z*_ at *s* = 0 (where **e**_*z*_ is a fixed basis vector in the lab frame) and **F** = **0, M** = **0** at *s* = *L*.

### A. Stable state at small m

Simulations are initialized from the vertically straight configuration over a range of values of *Sp, m*, and *k*. The filament quickly develops bending waves about a vertical time-averaged shape, or “backbone” (Fig. 2b, leftmost). If the forcing strength *m* is sufficiently small, or if *k* is negative (i.e. bending waves travelling from tip to base), the filament remains stable in this state.

### B. Whirling state at intermediate m

We henceforth focus on the mechanically more interesting regime of positive *k* (base-to-tip waves). If *m* is larger than some threshold, an instability emerges (Fig. 2b) in which a buckling deformation originates in the proximal region, and the filament settles into a whirling state consisting of fast, small-amplitude ‘minor’ waves superposed on the slowly whirling time-averaged backbone (also illustrated in Fig. 2c, top). Provided *k* and *Sp* are sufficiently high, whirling is attained for all *β*.

### C. Flapping states at large m

At larger values of *m*, a filament subject to circular helical forcing continues to exhibit whirling (albeit with reversed whirling direction and shape chirality). Under planar forcing, *β* = 0, a flapping state is instead attained (Fig. 2c, bottom) in which the backbone ‘flaps’ periodically in the plane normal to the minor waves, with the tip tracing a figure of eight. Near-planar forcing (*β* = *π/*16) produces flapping with slight whirling behavior: viewed from above, the backbone traces a slender ellipse rather than a straight line. This straight-whirling-flapping sequence with increasing *m* is reminiscent of an analogous transition in a filament under a follower force [40].

The two phase diagrams in Fig. 2(d-e) illustrate which of these states the filament evolves into depending on its mechanical parameters. At yet larger values of *m*, flapping itself becomes unstable; this occurs more readily at smaller *Sp*. The filament exhibits a whole zoo of behaviors, including chaotic beating, intermittent periodic planar beats, and temporary restabilization into whirling. A similar range of behaviors have also been reported in the tip follower force model in Ref. [45], which investigates larger forcing values than in Ref. [40]. Here, we do not classify these states in detail, instead simply regarding them as an unstable subset of the flapping solutions. We focus more on developing a detailed understanding of the first buckling instability leading to whirling.

### D. Instability emerges from compression

Next, we plot the time-averaged tension ⟨Λ⟩ in Fig. 2f for several values of (*m, k, Sp*). We find that ⟨Λ⟩ *<* 0 over virtually the entire filament, indicating that the filament is under compression. The observed transition may therefore be interpreted as a buckling instability. The fact that buckling first develops near the base is also consistent with the greater compressive tension in this region.

## IV. MULTIPLE-SCALES THEORY FOR A CLAMPED FILAMENT

We now perform a multiple-scales analysis to place this interpretation on a more rigorous footing and to determine how the rapid undulations generate the net compression responsible for buckling. The key here is the separation of timescales: the filament backbone evolves much more slowly than the fast minor undulations.

### A. Active internal moment generate fast travelling bending waves

We decompose the filament centerline as 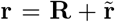 into a slow backbone **R** and a fast undulatory component 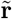 (see Fig. 2a), and further make the simplifying approximations that both the slow and fast components have small amplitudes, and that *k* is large. Balancing the 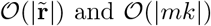 and *O*(|*mk*|) fast contributions in (1) yields the leading order equation for the fast dynamics, 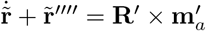. This has the particular solution

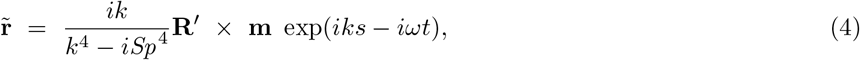

which describes travelling bending waves. This solution is valid in the bulk of the filament, but does not satisfy the boundary conditions at the ends. An additional standing wave complementary solution can remedy this but end effects may be safely neglected at this stage.

### B. Fast undulations rectify into mean compression

We now take the average of (2) over a period of fast undulation, using the fact the slow variable **R** remains approximately constant over a period. We again make a small-amplitude approximation for the slow filament shape, and only keep terms linear in the transverse slow filament shape **R**_⊥_.

Crucially, although 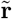 itself has zero time average, quadratic products of fast oscillatory terms have non-zero averages, thereby rectifying the fast oscillations into contributions to the slow balance. This rectification underlies the diverse filament dynamics thus far observed, and establishes them as genuinely nonlinear effects.

As detailed in SI Appendix, (2) time-averages to 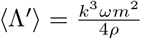, which may be integrated together with the force-free boundary condition, Λ = 0 at *s* = 1, to determine the time-averaged tension profile as

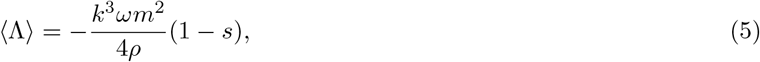

where *ρ* = *k*^8^ + *ω*^2^. For positive *k* and *ω* (base-to-tip flagellar waves), this is negative, and we have therefore derived the net compression responsible for the buckling instability. In Fig. 2f, we plot this theoretical result against the numerical mean tension profile, for various mechanical parameter values, showing excellent agreement; this also serves as justification for neglecting the end effects in the fast oscillatory solution. Our theoretical result also explains why filaments with tip-to-base waves remain stable: ⟨Λ⟩ is positive in this case, and the filament is stretched rather than compressed.

### C. Time-averaging full filament dynamics yields evolution equation for slow backbone

A similar time-averaging of (1) yields the following linearized equation for the slow dynamics,

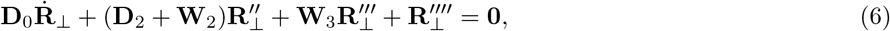

with the diagonal matrices 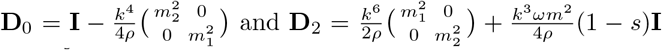, and the off-diagonal terms 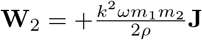 and 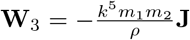, where **I** is the identity matrix and 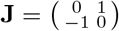. This equation characterizes the time evolution of the slow filament backbone and constitutes the central mathematical result of this study. Exploiting the separation of timescales, we have reduced the full filament dynamics described by (1) to explicitly determined fast undulations (4) superimposed on a slowly evolving backbone governed by the linearized equation (6). This is a substantial simplification of the strongly nonlinear three-dimensional dynamics, and provides a tractable framework for analyzing this buckling instability.

### D. Slow filament dynamics equation reduces to follower force model

Noting the similarities to classical follower-force-type equations [39, 40, 45], our result may be interpreted as a generalization of these equations. This comparison is made more precise by considering the limits of large sperm number and large wavenumber, specifically *ω* = *Sp*^4^ ≫ *k*^3^ and *k* ≫ 1, under which (6) reduces to

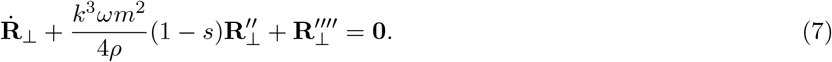

This is precisely the follower force equation, with a distributed tension equal to the profile derived in (5).

This result explains the similar sequence of stable, whirling, flapping, and chaotic states in our simulations and in follower-force studies [40, 45]. More broadly, it places the wave-induced buckling in Refs. [36, 37] and in the present study within the same framework as buckling driven by steady follower forces [39, 40, 45, 46]. The emergence of an effective follower-force equation from active moments distributed along the flagellum, naturally associated with dynein activity, also provides a concrete physical basis for a class of phenomenological models for ciliary and flagellar dynamics, whose direct physical motivation has previously been limited specifically to molecular cargo transport along cytoskeletal filaments.

### E. Linear stability of the slow dynamics reproduces the buckling threshold

We next perform a linear stability analysis of the slow equation to quantitatively determine the onset of buckling. Assuming normal mode solutions 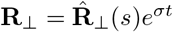, (6) yields an eigenvalue problem of the form 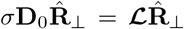, solved numerically by discretizing the operator ℒ using centered finite differences.

In Fig. 2g, we plot the real (solid) and imaginary (dotted) parts of the most unstable eigenvalue branches for different *β*, for the representative parameters (*Sp, k*) = (30, 8*π*). The most unstable branches are shown as thick curves, with the next most unstable branches shown as thinner curves in the same color scheme.

Here we have defined 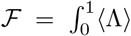 d*s* as the spatial mean of the time-averaged tension, allowing a convenient comparison with the classical problem of a filament subject to a follower force at its tip. In that case the tension is uniform and equal to the tip force, so that ℱ is equal to the applied tip force. For our undulating filament, we instead obtain ℱ = *k*^3^*ωm*^2^*/*8*ρ*. Owing to the similarities between follower force models and our slow equation, we recover eigenvalue spectra that are similar to earlier results for filaments subject to a follower force applied either at the tip or continuously along their length [39, 46, 47].

In the special case of a planar forcing, *m*_1_*m*_2_ = 0, the off-diagonal terms vanish, yielding two decoupled follower-force equations for the components *X* and *Y* of **R**_⊥_, each with slightly different tension terms, therefore giving rise to two nearly overlapping eigenvalue spectra (Fig. 2g, dark and light blue). The imaginary parts of the eigenvalues indicate a transition from monotonic to oscillatory stability at ℱ_1_ ≈ 18.02, while the real part of the leading eigenvalue becomes positive at ℱ_2_ ≈ 34.16, marking the onset of instability. These thresholds are similar to those of a filament subject to a tip follower force (ℱ_1_ ≈ 20.05 and ℱ_2_ ≈ 37.69) [39]. In the case of circular helical forcing, (6) can be rewritten as a single follower-force-type equation for the complex variable *W* = *X* + *iY*. The filament remains oscillatory stable until the onset of instability at ℱ ≈ 33.62.

The critical value of *m* at which the real part of the most unstable eigenvalue first becomes positive marks the instability threshold. In the phase diagrams in Fig. 2d-e, we compare this theoretical prediction (thick lines) with numerical stability outcomes and find excellent agreement, confirming that the observed buckling instability is governed by the derived slow filament dynamics.

### F. Unstable eigenmodes capture numerical filament shapes

In Fig. 2h, we also plot the eigenfunctions associated with the two most unstable eigenvalues. Since these are generally complex, their real parts represent the physical filament shapes. To illustrate the full envelope of each mode, we additionally superpose the real parts of *e*^*iϕ*^-multiples of the eigenfunction (dotted lines), with *ϕ* spanning [0, 2*π*].

A convincing validation of linear stability comes from the most unstable eigenmodes in the cases of planar and nearly planar (elliptical helical) forcing. The most unstable mode lies perpendicular to the beating plane in the planar case (Fig. 2h; left) and has a strong component perpendicular to the dominant beating direction in the near-planar case, consistent with out-of-plane disturbances exhibiting greater instability in simulations. The next most unstable eigenmode lies within the beating plane, and excitation of both modes with a phase difference produces whirling [40]. In contrast, in the stable flapping state, characterized by slow filament motion normal to the plane of the fast oscillations, it appears that only the most unstable mode is present.

The leading eigenmode for *β* = *π/*4 has a right-handed filament shape (Fig. 2h, right), consistent with the chirality observed in the simulations at moderate *m*. However, the eigenvalue branch of the next most unstable mode, shown by the light-red line, closely follows the leading branch, and at larger values of *m*, it is the sub-leading eigenmode that appears to be excited.

### G. Filament curvature and drag anisotropy convert undulations to mean compression

We have confirmed computationally that a filament can buckle under its own internal activity, and understood this mathematically as a result of the fast undulations rectifying into a mean compression. We now seek to understand the physical mechanism which converts these fast oscillations into a mean compression.

(2) may be understood as a tangential force balance on an infinitesimal section of the filament. Our calculations derived ⟨−**M**·**m**_*a*_⟩ = *k*^3^*ωm*^2^*/*2*ρ* and 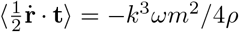, with the second term −**M**·**M**^*′*^ time-averaging to higher order (see SI Appendix). This may be interpreted as follows: the internal activity generates a tangential force, which the fluid drag partially balances, and the remainder appears as an internal compression.

The active term may be rewritten as −**M**·**m**_*a*_ = − (**t** ×**m**_*a*_)^*′*^ ·**t**, where **t** ×**m**_*a*_ is the transverse internal force induced by the active moment density. A tangential force density is then induced because this transverse force acts on a *curved* filament. This active term represents the tangential component of the change in a *transverse* active force along an infinitesimally small filament section, which is non-zero precisely because the direction transverse to the filament varies along the curved centerline (as clarified in the schematic in Fig. 2i).

For a more general drag anisotropy ratio *c*_⊥_*/c*_∥_ = *γ*, the hydrodynamic contribution is 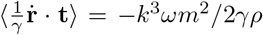. Under isotropic drag (*γ* = 1), tangential drag is strong enough to cancel the active compression. Such a filament experiences no net compression (⟨Λ⟩ = 0) and would not buckle through this mechanism. We have thus identified the mean compression as a consequence of activity along a curved filament, which tangential fluid drag only partially relieves in a medium with anisotropic drag.

## V. NUMERICAL SIMULATIONS OF A SWIMMING SPERMATOZOON

Although the clamped filament does not typically represent a biologically realistic setting, it was important to study because this isolates the buckling to its simplest setting, without the complications from a moving cell. Connecting this buckling to follower forces and establishing a concrete physical basis for these phenomenological models was also of broader importance. Equipped with a mathematical and physical understanding of the buckling in its simplest form, we are now ready to reintroduce swimming and return to our original biological motivation.

### A. Buckling instability in a swimming spermatozoon produces double waves

We extend our simulations to a freely swimming spermatozoon. Because insect sperm and many of the other species considered here have slender heads with widths comparable to that of the flagellum, we model the head by setting the region 0 *< s < ℓ* to have zero active moment and a bending rigidity 100 times that of the flagellar section (results are insensitive to the precise factor). The boundary conditions are now **F** = **M** = **0** at both *s* = 0 and *s* = 1. We present results for *ℓ* = 0.14, representative of the head-to-total-length ratio in many species; results are robust to moderate variations in *ℓ*.

Once the instability emerges, a filament under circular helical forcing propagates a large-amplitude helical major wave, with minor waves travelling along this backbone (Fig. 3a). The initially helical swimming trajectory is promoted to a superhelix after buckling. This important numerical result illustrates that the superhelical waveform can emerge purely as a mechanical instability. Although we focus on a representative *k* = 14*π*, double waves generically emerge for more general wavenumbers.

**FIG. 3.**
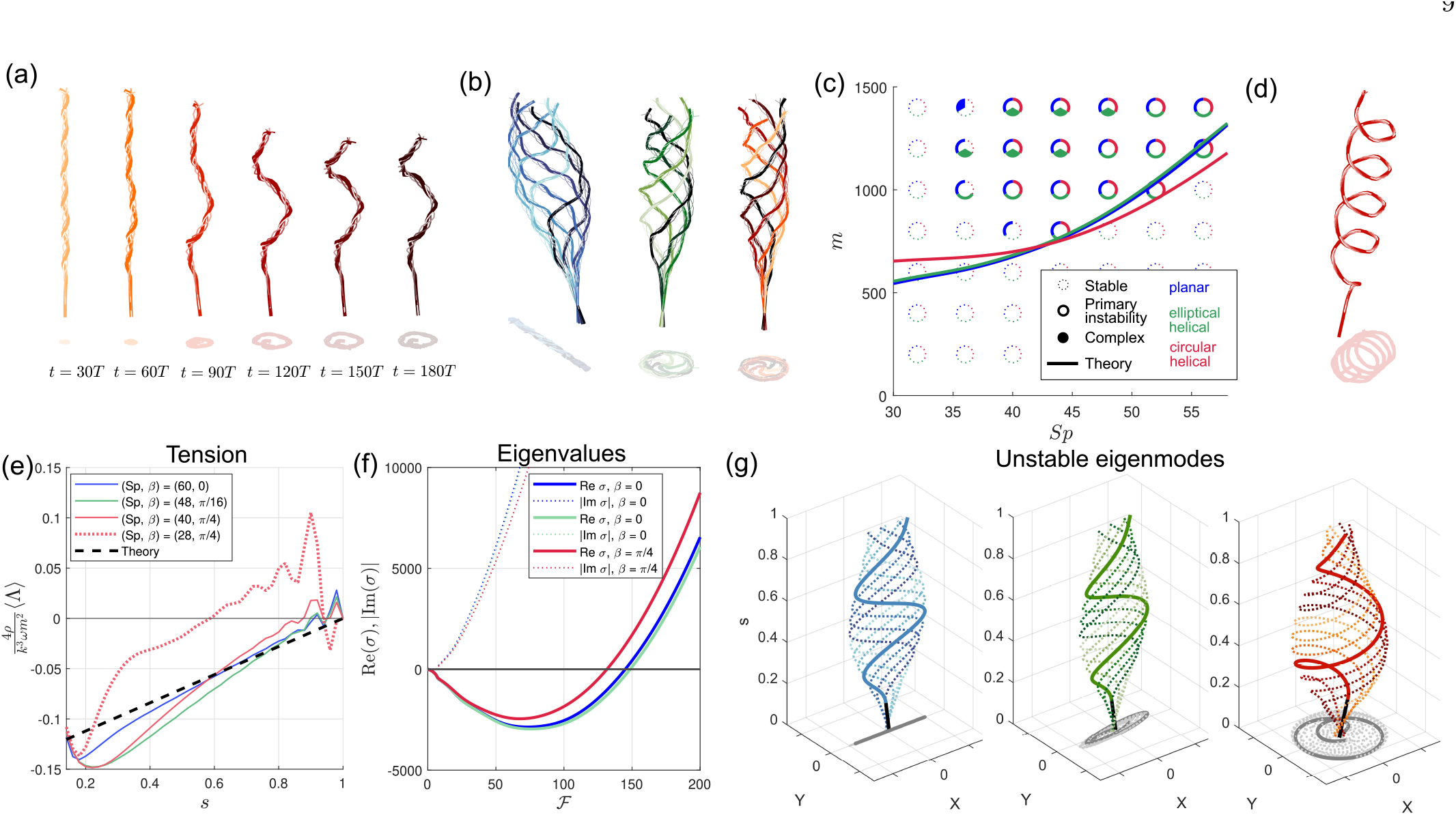
Numerical and theoretical results for a swimming spermatozoon. (a) Time sequence of a swimming spermatozoon, in the swimmer’s frame, buckling into a superhelical state. Filament snapshots from the preceding period are shown as thin lines to illustrate minor waves. Filament shadow is also shown. Parameters used are (*Sp, k, m, β*) = (44, 14*π*, 1400, *π/*4). (b) Filament snapshots at different times superposed to illustrate the planar double-wave morphology (left, *β* = 0) and superhelical morphology (center, right; *β* = *π/*16, *π/*4 respectively). Parameters used are (*Sp, k, m*) = (48, 14*π*, 1200). Time progresses from light to dark color. (c) Phase diagram illustrating long-time behavior for fixed *k* = 14 with (*m, Sp*) varied. Unfilled markers with dotted outlines indicate stable straight-swimming. Unfilled markers with solid outline indicates the primary instability illustrated in (b). Filled markers indicate more complex dynamics. *β* = 0, *π/*16, *π/*4 are indicated by blue, green, and red, respectively. Solid lines plot the critical buckling threshold *m*_*c*_ from linear stability analysis of the slow equation. (d) A spermatozoon buckled into a superhelical state with several major wavelengths, at large values of *k* and *Sp*, specifically, (*Sp, k, m, β*) = (200, 50*π*, 20000, *π/*4).(e) Time-averaged tension (normalized) computed numerically for various parameter values, plotted as colored lines, together with the theoretically calculated tension profile plotted as black dashed line. (f) Real (solid line) and imaginary (dotted) parts of the most unstable eigenvalue branch plotted against 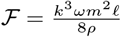, for (*Sp, k*) = (48, 14*π*). (g) The corresponding most unstable eigenmodes for *β* = 0 (blue), *π/*16 (green), and *π/*4 (red)

### B. Superhelical waveform is robust to ellipticity of active internal moment

Interestingly, a purely planar active moment (*β* = 0) develops a buckled state in which the major wave is planar and lies perpendicular to the plane of the rapid minor waves (Fig. 3b, left). We do not, however, expect this state to arise generically in nature, since real axonemal beating is unlikely to be perfectly planar. Introducing even a small degree of nonplanarity (*β* = *π/*16) recovers a buckled state with a helical time-averaged shape (Fig. 3b, middle), sharing the same time-averaged state with a circular helically forced filament (Fig. 3b, rightmost panel). This suggests that a helical major wave is the generic outcome of the instability regardless of the ellipticity of the active forcing, providing a natural explanation for the prevalence of superhelical waveforms across large variations in flagellar architecture.

### C. Secondary instabilities produce even richer dynamics

At large enough *m*, the filament may exhibit more complex dynamics, as indicated in the phase diagram in Fig. 3c. For the elliptical helical forcing, the superhelical swimming trajectory itself develops a larger-scale, coiled path. In the case of planar forcing, the angular amplitude of the major wave becomes so large that the spermatozoon cartwheels.

At substantially larger wavenumbers and sperm numbers, such as those attained by *Drosophila melanogaster* sperm which can reach lengths of up to 2 mm and accommodate as many as 70 minor waves [7], the filament can buckle into several major waves. Fig. 3d shows an example with (*k, Sp*) = (50*π*, 200). At large enough *m*, these major waves can themselves destabilize, producing a “super-superhelix.” We also observed states in which buckling is proximally confined, with a distal portion unbuckled and under extension. Although we focus here on the primary instability leading to a superhelix, these observations point to even richer dynamics in the largely unexplored regions of parameter space.

### D. Superhelical waveforms are restricted to large *Sp*

A less exotic, but arguably biologically more important observation is that, for a given *k*, buckling does not develop below a certain threshold of *Sp*. As we discuss later, this may provide an explanation for why some spermatozoa can sustain several helical waves without buckling, whereas longer spermatozoa with a similar number of minor waves buckle. What range of *Sp* permits buckling, and whether real spermatozoa have mechanical parameters within this regime, are important questions we will return to.

## VI. MULTIPLE-SCALES THEORY FOR A SWIMMING SPERMATOZOON

### A. Fast undulations generate mean compression

In order to better understand our numerical results, we now carry out the corresponding theoretical analysis. We work in the lab frame and make the approximation that the sperm swims in a straight line, say, with a velocity −*U* **e**_*z*_. In our time-averaging we also assume that the filament basis vectors remain approximately constant over a period of fast undulation, neglecting the small amount of rotation from the rolling of the cell about its swimming axis.

These calculations incorporate the swimming dynamics but otherwise remain largely similar to the clamped case, as detailed in SI Appendix, and yield the tension profile

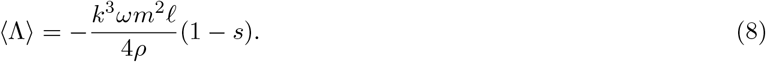

As in the clamped case, the fast undulations rectify into a net compression, thus confirming that the same buckling instability is at play here. Since the filament is now free to move against the viscous load of the head, rather than being firmly clamped, part of the compression is released, reducing it by a factor of *ℓ* relative to the clamped case. Provided *Sp* is high enough, our theoretical tension profile, plotted in Fig. 3e, agrees closely with numerical results.

### B. Linear stability of slow equation reproduces buckling threshold

To further quantify the resulting instability, we may carry out similar calculations time-averaging (1), yielding an equation characterizing the slow dynamics of the swimming spermatozoon. This equation is similar to (6) for the clamped case, and its precise form is detailed in SI Appendix. A linear stability analysis recovers its eigenvalues (Fig. 3f) and reveals a stability threshold *m*_*c*_ in excellent agreement (at large *Sp*) with simulations (Fig. 3c), confirming our picture of buckling driven by mean compression induced by the fast undulations.

### C. Unstable eigenmodes broadly capture numerical waveforms

The most unstable eigenmode (Fig. 3g, right) in the case of circular helical driving is a right-handed circular helix, consistent with simulation results. The dominant eigenmode is a right-handed elliptical helix in the case of an elliptical helical active moment (Fig. 3g, center). Under nonlinear interactions and rolling of the cell, this mode evolves into a superhelix with a roughly circular helical major wave. In the planar case (Fig. 3g, left), simulations and the most unstable eigenmode both exhibit buckling confined to the plane normal to the fast oscillations.

Although not biologically relevant for the long spermatozoa that exhibit double waves, we find that increasing the filament radius from *ϵ* = 0.001 to 0.01 reverses the major wave chirality, overturning the naive expectation that aspect ratio should have no qualitative effect on the dynamics. A filament driven by right-handed fast bending waves thus buckles into a left-handed helical backbone at *ϵ* = 0.01. This surprising numerical result arises from the hydrodynamic torque on the filament, which we had neglected because it is quadratically small in *ϵ*. Reincorporating this torque and filament twist, as detailed in SI Appendix, recovers eigenmodes with the chiralities observed in simulations. The unexpected behavior at larger *ϵ* provides a cautionary example of how an asymptotically and numerically small hydrodynamic torque can nevertheless have a decisive qualitative effect.

### D. Decreasing *Sp* lowers proportion of filament under compression

Although the theoretical tension profile closely matches simulations at large *Sp*, a discrepancy is seen at lower *Sp* (Fig. 3e, dotted red). Since compression causes buckling, this discrepancy has direct implications on linear stability. As *Sp* decreases, the theoretical *m*_*c*_ decreases, failing to capture the sharp increase in the numerical threshold (Fig. 3c). We seek to reconcile these differences.

Our theoretical solution thus far, which we recall neglects end effects, correctly reproduces the slope of the time-averaged tension profile in the bulk of the filament, but does not capture the sharp variation of ⟨Λ⟩ in the boundary layers at the proximal and distal ends. This becomes more pronounced as *Sp* decreases, and effectively shifts the tension profile upwards, thus decreasing the portion of the filament under compression (⟨Λ⟩ *<* 0). At *Sp* = 28, only approximately half of the filament is under compression (Fig. 3e, dotted red), in stark disagreement with the theoretical approximation predicting compression in the entirety of the filament (Fig. 3e, dashed black). A shorter portion under compression makes compression-driven buckling more difficult to attain, and thus requires higher values of the active forcing for instability.

### E. End corrections reconcile discrepancies at lower *Sp*

To further quantify these findings, we incorporate end corrections into our theoretical analysis, summarised here and detailed in SI Appendix. As discussed in §IV A, the fast undulations comprise a travelling wave particular solution (which dominates the bulk of the filament at large *Sp*), and a standing wave complementary solution thus far neglected. This standing wave solution is 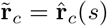 exp(− *iωt*), where 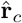 satisfies 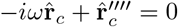 and is therefore a linear combination of complex exponentials with growth/decay rates of *ω*^1*/*4^ = *Sp*. This indicates boundary layers at the filament ends of width *O*(1*/Sp*), explaining the larger influence of end effects at lower *Sp*.

As illustrated in Fig. 4a, the travelling wave dominates at high *Sp* (left panel), with standing wave behavior confined distally, whereas the standing wave dominates at lower *Sp* (right). Our tension calculations thus far retain only the travelling wave, yielding a linear profile that is everywhere compressive. Reintroducing the standing wave produces a more tension profile in better agreement with simulations (Fig. 4b). This also provides additional validation of the numerical scheme in the present setting, complementing its extensive validation for other problems in Ref. [35]. We have thus quantified the end effects and identified them as the source of the discrepancy between the theoretical and numerical stability thresholds.

**FIG. 4.**
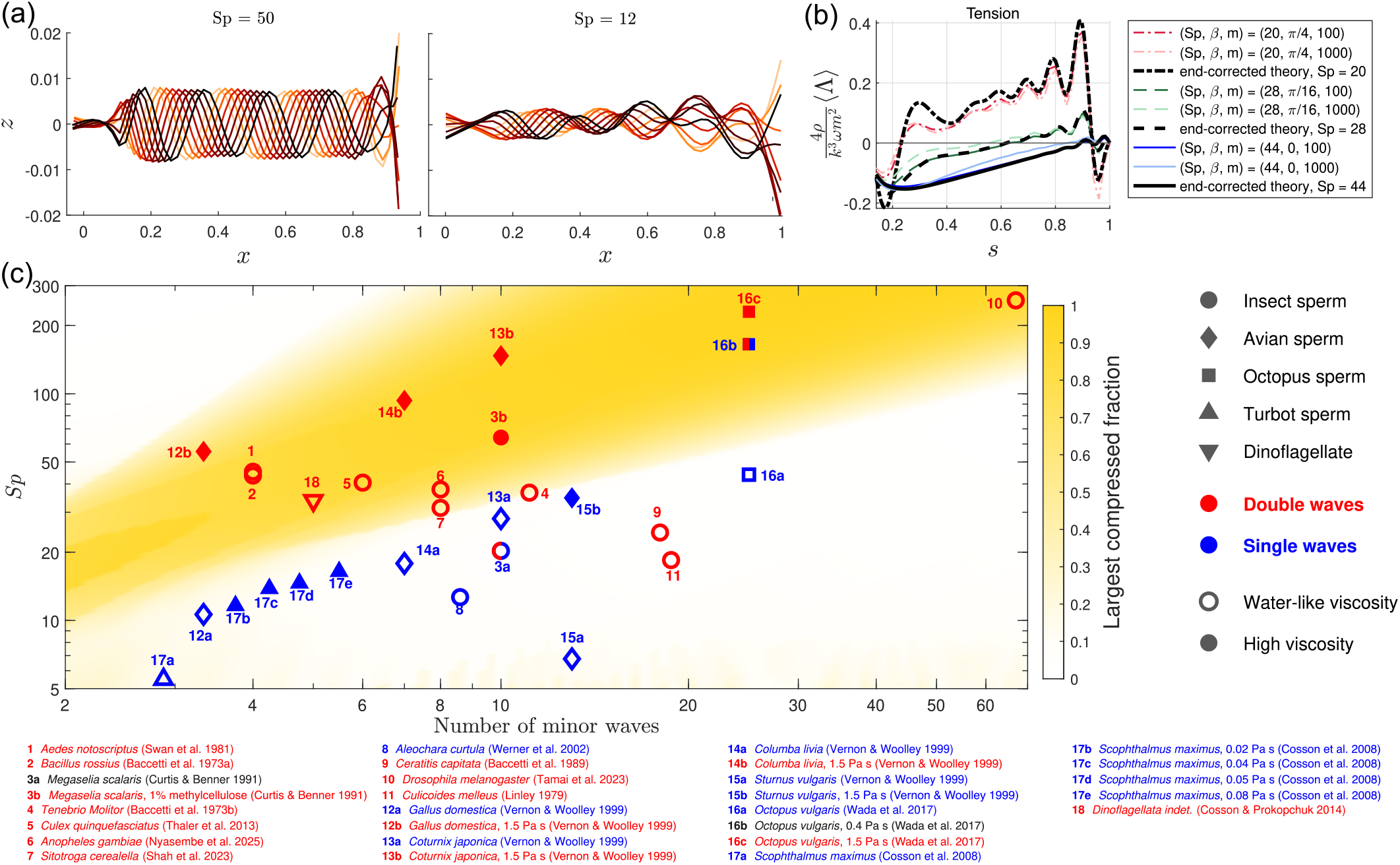
Comparison of buckling regime with biological parameter estimates. (a) Fast undulations over a period from simulations of a swimming spermatozoon, with time progressing from light to dark color, illustrating a dominant travelling wave behavior at high sperm number *Sp* = 50 (right) and a dominant stand wave behavior at low sperm number *Sp* = 12. Parameters used are *k* = 10*π* and *m* = 2000 (left), 100 (right). (b) Numerically computed time-averaged tension profile for various parameter values (color lines) plotted together with end-corrected theoretical solution (thick black line). (c) Heat map of largest compressed fraction of filament, in the parameter space of (*Sp, k*). Buckling is possible in the dark region (corresponding to a high largest compressed fraction). Parameter estimates of various species are overlaid, with red(blue) indicating flagella exhibiting double(single) waves. Different symbols indicate taxonomic diversity. Unfilled markers with solid outline indicate cells swimming in a fluid of water-like visosity, while filled markers indicate higher viscosity. Data points sharing the same number indicate experiments on the same species at different viscosities.

### F. Largest compressed fraction of the filament identifies the buckling regime

In the spirit of the preceding theoretical analysis, one may in principle derive an end-corrected version of the slow dynamics equation, and via a linear stability of said equation, determine the buckling threshold. Although this is straightforward, it is cumbersome and yields no additional physical insight. We instead see that the end-corrected tension profile already provides ample information about the possibility of buckling.

From the end-corrected theoretical tension profile, we may compute the proportion of the total filament length that the *largest* connected region under compression occupies, henceforth referred to as the largest compressed fraction. For instance, the distal ends of the numerical tension profiles in Fig. 3e and Fig. 4b contain a small compressed region, but this is disconnected from the larger compressed region in the filament bulk and therefore does not contribute to buckling. It is the *largest* compressed fraction that determines whether buckling is possible.

Due to finite computational resources, numerical instabilities at excessively large *m*, and the expansive three-dimensional (*Sp, k, m*) parameter space, it is impractical to determine, for each *k*, a strict threshold of *Sp* below which buckling cannot occur. Although the largest compressed fraction does not provide a strict mathematical threshold, it circumvents these difficulties and nonetheless gives a reliable indicator of the possibility of buckling. Fig. 4c shows how the largest compressed fraction varies with sperm number and the number of minor waves *N* (regarded interchangeably with wavenumber *k* = 2*πN/*(1 −*ℓ*)), with colors close to white indicating largest compressed fractions close to zero. Importantly, we observe a dark band within which buckling is possible.

## VII. COMPARISON WITH BIOLOGICAL OBSERVATIONS

We have shown conclusively that double waves can emerge purely as a mechanical instability from a filament subject to a single set of active bending waves. Whether double waves emerge in real biological spermatozoa via this mechanism, however, is a more nuanced issue.

### A. The buckling regime broadly separates double and single waves

We surveyed the literature for spermatozoa whose flagella exhibit helical (single-wave) or superhelical (double-wave) morphologies, and for each species estimated the sperm number *Sp* = *L*(*c*_⊥_*ω*_∗_*/EI*)^1*/*4^, and the number of minor wavelengths along the flagellum. Since measurements of flageller bending rigidities are unavailable, the measured order of magnitude for eukaryotic flagella, *EI* = 10^−21^ N m^2^ [48–51], is adopted. Apart from the sperm length *L*, the sperm number depends on the other physical quantities only as one-fourth powers, so uncertainties in their estimates do not mitigate strongly.

We populate Fig. 4c with (*k, Sp*) estimates for each species, indicating instances of double (single) waves in red (blue). The blue points include sperm from fish, birds, and octopuses. We omit spermatozoa with single helical waves of fewer than 2.5 turns, since a superhelix is difficult to distinguish with few minor turns. All instances of single waves lie in the nearly white region where buckling is prohibited, with the exception of octopus sperm in 0.4 Pa s methylcellulose solution. This is not inconsistent with theoretical predictions since the buckling regime permits but does not guarantee buckling; this may simply reflect forcing not strong enough to cause buckling. Double waves are observed as expected when ciliobrevin-D, which suppresss sea urchin sperm motility [52] but modifies octopus sperm dynamics less dramatically, is added.

We also see that the (*k, Sp*) values of most instances of double waves, spanning the sperm of insects, birds, and octopuses, and dinoflagellates, lie within the dark band where buckling is possible. The correspondence between the buckling regime and the occurrence of double waves is consistent with an elastohydrodynamic mechanism, although the few exceptions warrant further discussion.

### B. Exceptions in insect sperm are suggestive of mechanisms beyond elastohydrodynamics

The tephritid fly *Ceratitis capitata* [53] lies outside the buckling regime but exhibits double waves with several major waves. These spermatozoa can also swim backwards, and exhibit double waves during both forward and backward swimming. This appears to contradict our elastohydrodynamic explanation, which prohibits buckling in the backwards-swimming state where minor waves travel proximally. However, closer investigation is warranted to confirm that major waves are genuinely actively generated in the backwards-swimming state and not merely a remnant of the major waves produced during forward swimming. Nevertheless, the available evidence suggests a more complicated picture than a pure elastohydrodynamic instability for this species. *Tenebrio molitor* and *Sitotroga cereallela* also lie just within the buckling region, very close to its boundary, and warrant further investigation.

Interestingly, *Culex quinquefasciatus* sperm exhibit both double waves and long-wavelength helical single waves. Although the double waves have minor-wave count and sperm number consistent with buckling, this species provides an example in which the same flagellum can actively generate (although not necessarily simultaneously) bending waves of different wavelengths.

### C. Special cases complicate the classification of double waves

Double waves are also seen in the sperm of the biting midge *Culicoides melleus* [6], despite its (*k, Sp*) estimate predicting that buckling is inhibited. However, the major helix here is extremely slender (see Fig. 1h): with a major wave amplitude (mean 2.1 µm) merely a hundredth of the mean cell length [6]. Furthermore, these sperm were observed under the extreme comfinement of a fluid layer less than 4 µm thick. This study therefore presents very different conditions from our theory and simulations, developed in the simpler setting of an unbounded fluid.

Phenomena different from propagating major and minor waves have been grouped together with genuine double waves [10]. *Aleochara curtala* spermatozoa appear to exhibit double waves along its relatively short flagellum (Fig. 1f), but these are in fact single waves along an intrinsically curved flagellum which rolls with the cell [9]. Although this was proposed as an alternative explanation in the double waves debate, this mechanism may not generalize beyond this particular genus. We have indicated this in Fig. 4c as a species exhibiting single waves.

Ref. [8], altogether omitted from Fig. 4c, describes minor waves propagating along an already coiled centerline, with the large coils lying approximately in a single plane. This is a very different geometry from the genuinely three-dimensional superhelices produced by propagating major and minor waves. Barring these special cases, the numerous remaining studies surveyed clearly report genuine double-wave morphologies, in which both the major and minor waves propagate along the flagellum.

### D. *Megaselia scalaris* sperm indicate multiple mechanisms at play

The spermatozoa of the humpbacked fly *Megaselia scalaris* both illustrate an elastohydrodynamic mechanism as well as some less clear origin of double waves. The sperm cells display a “rounded” state in which minor waves can propagate along a flagellum coiled into a circle, as well as a forward-swimming superhelical state [5]. We interpret the rounded state as an unbuckled filament with intrinsic curvature and possibly subject to near-wall hydrodynamic or steric effects. Both rounded and superhelical cells are seen in a water-like medium, and these superhelical cells thus exhibit double waves outside the buckling regime. However, in a more viscous 1% methylcellulose solution, most cells become superhelical, consistent with a higher *Sp* promoting buckling. This is therefore a very interesting case possibly illustrating multiple effects at play.

### E. Viscosity-dependence of double waves supports an elastohydrodynamic mechanism

The avian and octopus spermatozoa, for which the same cells were observed at different viscosities (indicated in Fig. 4c as points sharing the same number label) and double waves appeared only in viscous media, provide strong evidence for an elastohydrodynamic mechanism. The spermatozoa of the domestic fowl, pigeon, and quail were observed to exhibit double waves in a highly viscous medium (1.5 Pa s methylcellulose solution), but not in a water-like medium. Although the authors attributed the superhelical waveforms to the elongated helical heads of the avian spermatozoa, this argument does not explain their absence at low viscosity. Our elastohydrodynamic mechanism not only reproduces the superhelical waveform, but also offers an explanation for why double waves only appear in high-viscosity media.

### F. Sequential emergence of minor and major waves supports a buckling mechanism

Beyond simply distinguishing between single or double waves, Ref. [3] reports the time progressions of these waves: small-amplitude minor waves are first observed, followed by emergence of the larger amplitude major waves. The sequential appearance of these waves is consistent with the onset of a buckling instability in a filament initially possessing only the minor waves.

## VIII. CONCLUSION

### A. Summary of results

Motivated by the recurrence of double waves in spermatozoa across diverse taxa and flagellar architectures, we hypothesized that this phenomenon may have a mechanical origin, specifically, that double waves can mechanically emerge from a filament exhibiting single waves. Using numerical simulations and a theoretical analysis, we have shown that a buckling instability indeed generates double waves in an actively undulating filament.

We first isolated this mechanism in the simpler setting of a clamped filament. Averaging over the active undulations yielded an equation for the slow filament dynamics, which reduces to the follower-force model. Linear stability accurately predicted the critical active moment strength above which buckling occurs, while the unstable eigenmodes corresponded closely to the dynamics seen in simulations. We further identified the physical origin of the instability as an effective compression, generated by active bending moments acting along a curved filament in a fluid medium with anisotropic drag.

We then returned to our original biological motivation and extended the numerical and theoretical analysis to a swimming spermatozoon, showing that the same buckling mechanism produces superhelical waveforms. Incorporating end corrections enabled quantitative prediction of the tension profiles, allowing us to characterize the mechanical regime within which buckling can occur.

### B. Discussion

Comparison with the biological literature supports an elastohydrodynamic mechanism. Double waves recur in spermatozoa across diverse taxa and flagellar architectures, and our simulations reproduce these waveforms purely mechanically. Parameter estimates for numerous species place most reported double waves within the regime in which our theory allows buckling. The remarkable diversity of sperm structure makes it difficult, without highly targeted experiments in individual species, to completely rule out additional biological mechanisms underlying double waves. Nonetheless, their recurrence across diverse taxa, together with their consistency with the buckling regime, strongly suggests that elastohydrodynamic buckling plays a pervasive role in their emergence.

These results also highlight the importance of the physical environment in interpreting biological observations. It remains unknown whether such superhelical waveforms persist in nature, where large numbers of spermatozoa interact within the strong confinement of reproductive tracts. The emergence of double waves may depend sensitively on factors such as viscosity and confinement, yet in our literature survey we encountered many studies in which viscosity was not measured or reported. Similarly, the mechanical environment experienced by sperm cells was rarely described. Our model assumes an unbounded fluid, a reasonable approximation for a sufficiently deep microscope chamber, but in an overly thin fluid layer, hydrodynamic confinement introduces an additional level of complexity which complicates the interpretation of observed waveforms. Closer interaction between the life and physical sciences may therefore help identify variables which may appear secondary from the perspective of one discipline but become essential in a more holistic approach.

In our endeavor to address a biological question, we have also answered questions in the mechanics of active elastic filaments. Although prior simulations have reported similar instabilities [36, 37], our multiple-scales treatment has analytically quantified the activity-induced compression and identified the roles of internal activity, curvature, and fluid drag, thus providing a more complete understanding of the physical mechanism underlying this nonlinear buckling. The slow dynamics equation emerging from the undulations places this phenomenon within the framework of classical follower-force instabilities, thereby lending greater biological relevance to, and providing a concrete physical basis for, this important class of phenomenological models for active biological filaments.

There are several natural extensions of the present theory. We have used resistive-force theory, thus accounting only for the dominant local hydrodynamics. It would be interesting to investigate how nonlocal hydrodynamic interactions [54–56] between distant parts of the flagellum modify the instability threshold and the nonlinear waveforms. Incorporating confinement presents another challenging but important extension, because it would more faithfully reproduce *in vitro* experimental geometries and the strong confinement in reproductive tracts. Relaxing the assumption of an isotropic filament [57] could more realistically model certain spermatozoa. Nevertheless, the success of our minimal model in generating a buckling instability which qualitatively reproduces the experimentally observed double waves suggests that it captures the fundamental physical mechanism.

Our study expands the study of flagellar motility beyond conventional model organisms, and shows that similar superhelical waveforms can arise in evolutionarily distant taxa through a common elastohydrodynamic principle. Our results have implications for how flagellar mechanics can shape sperm responses to the different environments encountered during reproductive transport and fertilization. These results may also inform the design of artificial microswimmers, in which self-buckling could be exploited to achieve complex gaits and trajectories, or suppressed when large-scale instability is undesirable. More broadly, our findings illustrate how complex, highly organized biological patterns can emerge from small-scale activity and mechanical instability.

## MATERIALS AND METHODS

### C. Numerical scheme

The elastohydrodynamic equations are solved using the computational scheme of Ref. [35], which extends the asymptotic coarse-grained approach of Ref. [34] from planar to three-dimensional filaments. This requires repeated changes of basis to avoid coordinate singularities. Spatial derivatives are discretized using fourth-order finite differences, and the resulting system is integrated in time with MATLAB’s stiff ODE solver `ode15s` [58]. We discretize the filament into *N* = 50 segments, with the exception of Fig. 3d, which uses *N* = 150.

### D. Octopus spermatozoa experiments

Observations of *Octopus vulgaris* sperm were previously presented by Yuuko Wada and Shinji Kamimura at the 22nd International Congress of Zoology and the 87th Meeting of the Zoological Society of Japan (Okinawa, Japan, 2016). We provide the experimental details relevant to the data used here. Formal animal ethics approval was not required for experiments involving cephalopods under Japanese regulations. Adult *Octopus vulgaris* were dissected under cold anesthesia and spermatozoa were collected from the Needham’s sac (male) or from near the spermathecae (female). Spermatozoa were diluted in artificial seawater containing 0.1% BSA. High-viscosity media contained 2% (w/v) methylcellulose (0.4 Pa s or 1.5 Pa s). Where indicated, 100 µM ciliobrevin-D was added. The observation chamber depth was set using 58 µm-thick Scotch mending tape as a spacer. Sperm motility was observed by phase-contrast microscopy using a 10× objective and recorded for 1 s at 200 fps using a HAS-220 high-speed CCD camera, or for 10 s at 30 fps using an iDS UI-3240CP camera.

### E. Biological parameter estimates

Since direct measurements of flagellar bending stiffnesses are unavailable for the species considered here, we use *EI* = 10^−21^N m^2^, corresponding to an order-of-magnitude value for the flagella of sea urchin sperm [48–50] and of *Chlamydonomas* [51], another model organism for flagellar propulsion. The minor wave frequency is often reported, but when measurements are not available we use a value of 30 Hz as a reasonable order-of-magnitude estimate. Many of the studies surveyed use dilute aqueous salt solutions, whose viscosities are expected to be approximately that of water, and we therefore use *µ* = 10^−3^ Pa s in our estimates, except when a high-viscosity medium is specifically indicated. Further details are included in SI Appendix.

## ACKNOWLEDGEMENTS

We are grateful to Yuuko Wada (Ochanomizu University) for performing the octopus spermatozoa experiments and generously providing the data. P. H. H. and K. I. acknowledge the Japan Society for the Promotion of Science (JSPS) KAKENHI for JSPS International Research Fellow (Grant No. 25KF0222). K. I. acknowledges JSPS KAKENHI (Grant No. 24K21517) and the Japan Science and Technology Agency (JST), FOREST (Grant No. JPMJFR212N) and CREST (Grant No. JPMJCR25Q1).

## Notes

### Competing Interest Statement

The authors have declared no competing interest.

